# Full Scan enhanced Dynamic Range MS improves metabolite coverage and cancer cell-line discrimination in untargeted metabolomics

**DOI:** 10.64898/2026.06.11.731534

**Authors:** Demi J. Rijlaarsdam, Michał Kaczmarek, Christian Klaas, Christian Thoeing, Kyle L. Fort, Susan S. Bird, Celia R. Berkers, Esther A. Zaal

## Abstract

Metabolite detection with mass spectrometry (MS) in untargeted metabolomics is limited by the wide concentration range of metabolites, where high-abundance signals dominate MS^1^ scans and suppress detection of low-abundance features. This reduces metabolite coverage and obscures biologically relevant signals, particularly in complex cellular systems. Full Scan enhanced Dynamic Range (eDR) MS addresses these limitations by partitioning the MS^1^ mass range into multiple subscans and mass windows, reducing saturation effects from dominant ions. Here, we systematically evaluate different eDR acquisition strategies for untargeted metabolomics.

Across four hepatocellular carcinoma cell lines, Full Scan eDR MS increased detectable features up to ∼3.5-fold compared to Full Scan MS. Among equidistant window configurations, 12 windows yielded the highest feature count and broadest dynamic range, while custom window distributions further improved detection in ion-dense regions. In particular, allocating smaller window sizes to the low m/z region selectively increased detection of low-mass features while preserving performance for higher mass ions. Full Scan eDR MS also improved data quality, reducing variation and increasing signal-to-noise ratios, especially for low-abundance metabolites. MS^2^ coverage and metabolite identifications increased substantially, resulting in unique detection of cancer-relevant metabolites. Importantly, the increased depth of metabolite detection enabled improved discrimination between cancer cell lines, supporting deeper interrogation of metabolic heterogeneity.

Overall, these results establish Full Scan eDR MS as a flexible strategy to improve sensitivity and metabolome coverage in untargeted metabolomics. Customization of window size and distribution enable targeted expansion of dynamic range within predefined mass regions, allowing MS acquisition to be tailored to sample complexity and metabolites of interest.

## Introduction

Cellular metabolism integrates genetic and environmental influences into functional outcomes, thereby representing the molecular layer closest to the phenotype. Consequently, metabolomics, the comprehensive profiling of metabolites, is essential to understand biological systems and phenotypes. Metabolomics is widely applied across biological and biomedical research, including plant, microbial and mammalian systems (Alseekh et al., 2021; Fernie et al., 2004; Rinschen et al., 2019). In cancer research, metabolomics has become an essential tool for studying the metabolic reprogramming that drives tumor development and progression (Hanahan, 2026; Pavlova et al., 2022), and identifying novel metabolic targets (Cai et al., 2025; Liu et al., 2022; Mishra et al., 2025). The rapid expansion of metabolomics studies reflects the need for advanced analytical strategies capable of capturing metabolites associated with complex biological processes and diseases, including cancer, aging, inflammation and neurodegenerative diseases (Huang et al., 2025; Moreau et al., 2020; Shao et al., 2021; Sun et al., 2024). These advances have also facilitated the application of metabolomics in clinical research and biomarker discovery (Kennedy et al., 2018; Sun et al., 2024).

Liquid chromatography-mass spectrometry (LC-MS)-based metabolomics has emerged as a key analytical platform for the detection and quantification of metabolites across diverse biological systems (Furlani et al., 2021; Guijas et al., 2018; Shao et al., 2021) in both targeted and untargeted approaches. Untargeted metabolomics aims to capture a comprehensive and unbiased snapshot of all detectable metabolites within a sample (Schrimpe-Rutledge et al., 2016; Souza & Patti, 2021). This approach not only results in the broadest possible coverage of the metabolome, but has also proven powerful for discovering novel metabolites (Gelambi & Whitehead, 2024; González-Domínguez et al., 2023; Kharazian et al., 2025; X. Wu et al., 2025). Modern LC-MS based untargeted workflows typically combine high-resolution MS^1^ acquisition with data-dependent MS^2^ (ddMS^2^) fragmentation scans (Pezzatti et al., 2020; Y. Wu & Wang, 2025). Recent advances, such as iterative exclusion approaches like AcquireX intelligent data acquisition, have been implemented in instrument platforms to enhance metabolome coverage by systematically reducing redundant fragmentation events and prioritizing previously unfragmented features (Cooper & Yang, 2024; David et al., 2021).

Despite these advances, achieving high sensitivity, broad metabolite coverage and confident metabolite identifications remain a major challenge in untargeted metabolomics (David et al., 2021; Schrimpe-Rutledge et al., 2016; Shen et al., 2019; Theodoridis et al., 2026). A key limitation is the wide concentration range of metabolites, where high-abundance ions dominate MS^1^ scans and suppress the detection of low- and medium-abundance compounds in complex matrices (Cui et al., 2018; Lu et al., 2017; Meister et al., 2021). Strategies that segment the mass range into narrower m/z windows, such as BoxCar and spectral stitching approaches, have been shown to improve signal-to-noise ratios and enhance detection of low-abundance molecules by reducing ion saturation of high abundance species (Kaufmann et al., 2020; Meier et al., 2018; Ranninger et al., 2016; Sarvin et al., 2020; Southam et al., 2017). However, these segmented acquisition strategies result in multiple partial MS^1^ spectra, requiring nontrivial post-processing and calibration steps to reconstruct the Full MS spectrum and correct the signal intensities at the edges of the isolated m/z windows. Also, the lack of available Full MS spectra during acquisition limits the online data-dependent capabilities of the instrument.

To overcome these dynamic-range limitations, a new acquisition strategy was developed, termed Full Scan enhanced Dynamic Range (eDR) MS, by implementing intelligent and automated MS^1^ multiplexing strategies with optimized injection times. This resulted in the novel high resolution mass spectrometry (HRMS) eDR scanning mode on the Thermo Scientific™ Orbitrap™ Excedion™ Pro Mass Spectrometer, which increases metabolite coverage in complex samples. In eDR acquisition, the full MS^1^ mass range is divided into two Orbitrap subscans, each encompassing multiple m/z windows that are non-sequentially isolated by the quadrupole and accumulated into a single Orbitrap injection. The two Orbitrap subscans are then seamlessly integrated into a single multiplexed MS^1^ full spectrum (Figure 1). By splitting the mass range and assigning individual ion injection times based on the ion flux per isolation segment, this approach enables the detection of both low- and high-abundance compounds with improved signal-to-noise ratio (S/N), measuring a higher dynamic range in complex samples. Moreover, flexible customization of window numbers and distribution enables targeted optimization of metabolite detection within specific m/z regions.

**Figure 1.**
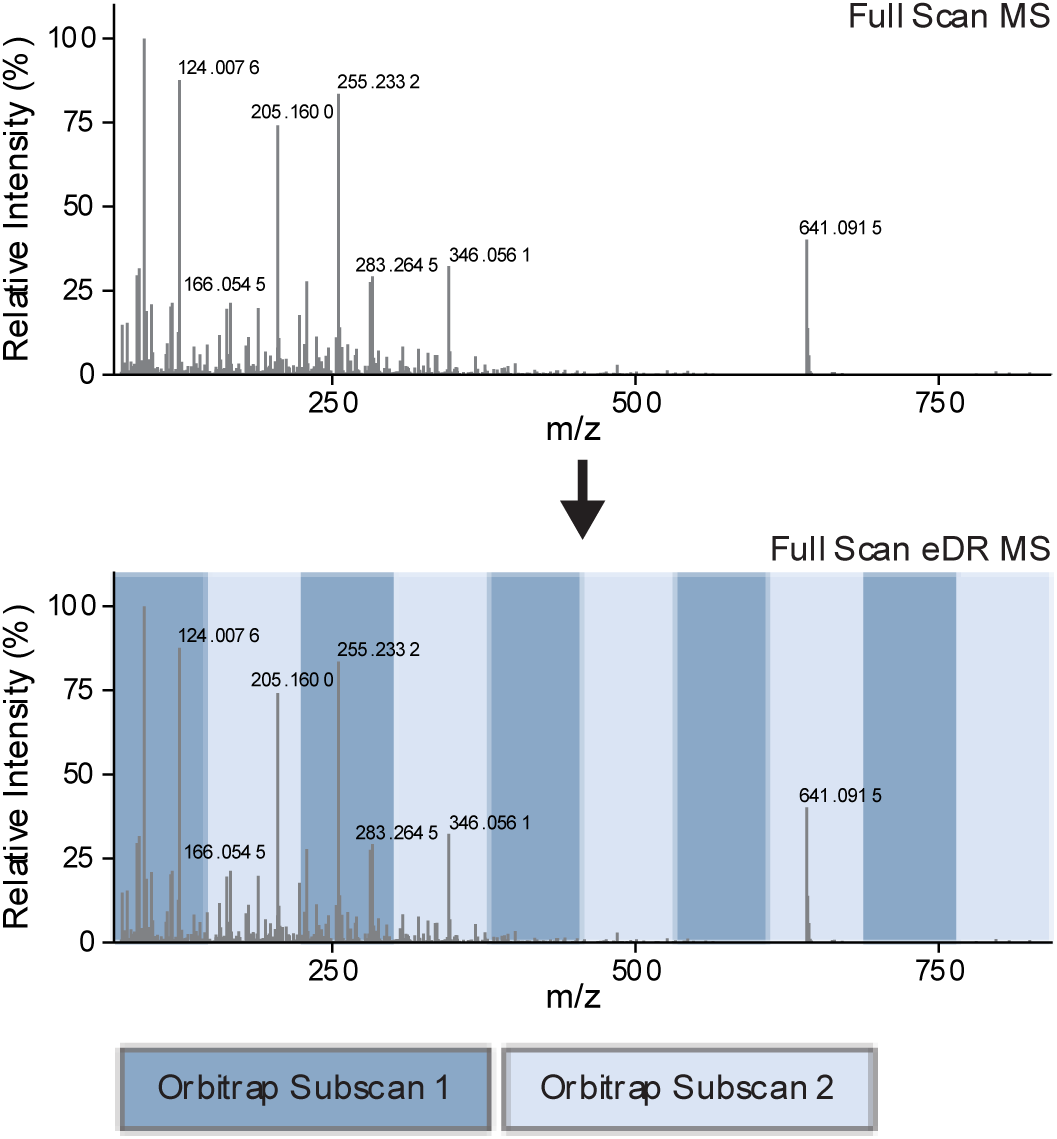
eDR overview. Schematic representation of window segmentation in Full Scan eDR MS acquisition. In conventional Full Scan MS acquisition, ions across the entire selected m/z range are accumulated and detected together in a single Orbitrap scan (top). In eDR mode, the m/z range is segmented into alternating isolation windows that are acquired in two separate Orbitrap subscans (bottom). Within each subscan, the quadrupole non-sequentially isolates the predefined m/z windows, and the corresponding ions are collected and detected together in the Orbitrap within a single injection. The two subscans are then combined into a single multiplexed MS^1^ spectrum, reducing ion competition and improving detection of low-abundance metabolites.

In this study, we systematically evaluate the performance of eDR acquisition for untargeted metabolomics across multiple hepatocellular carcinoma (HCC) cell models. HCC is characterized by extensive metabolic rewiring that supports tumor growth and progression, including alterations in glycolysis, the pentose phosphate pathway (PPP) and the tricarboxylic acid (TCA) cycle (Du et al., 2022; Park & Hall, 2025). Enhanced glycolytic flux despite oxygen availability, commonly referred to as the Warburg effect, represents one of the best-known examples of these metabolic adaptations (Vander Heiden et al., 2009). As metabolic dysregulation is a defining feature of HCC progression, comprehensive metabolomic profiling is essential for understanding disease-associated metabolic heterogeneity.

Using this model, we compared multiple eDR window configurations and distributions to determine their impact on metabolite detection and identification. We demonstrate that eDR improves metabolite coverage, enhances data quality and enables detection of biologically relevant metabolites in HCC that remain inaccessible under conventional untargeted acquisition methods.

## Materials and methods

### Cell lines and culture conditions

Human hepatocellular carcinoma (HCC) cell lines HepG2, Hep3B, Huh-7 and SNU-449 were obtained from ATCC. Cells were cultured in William’s E Medium (Thermo Fisher Scientific, #A12176), supplemented with 10% fetal bovine serum (FBS, Thermo Fisher Scientific, #A5256701) and 1X GlutaMAX (Thermo Fisher Scientific, #35050061). Cells were cultured at 37°C in humidified atmosphere containing 5% CO_2_. All cell lines were tested for mycoplasma contamination every 2 months.

### Cell Lysates

Cells were seeded in T75 flasks for 3-4 days to reach ∼90% confluency. For harvesting, media was aspirated and cells were washed with ice-cold 1X Phosphate-Buffered Saline (PBS). Cells were detached using 0.05% Trypsin-EDTA (1X) (Thermo Fisher Scientific, #25300), after which cell culture medium was added to neutralize trypsin. Cells were collected in 50 mL tubes, centrifuged at 500 x g for 5 minutes at 4°C, washed twice with ice-cold PBS, resuspended in PBS and 1×10^6^ cells were transferred to cold Eppendorf tubes. Cells were centrifuged at 500 x g for 5 minutes at 4°C, supernatants were discarded and cell pellets were snap-frozen in liquid nitrogen. Pellets were stored at −80°C until LC-MS analysis. Frozen cell pellets were processed immediately prior to LC-MS measurements to ensure fresh lysates. For metabolite extraction, 200 µL of MS lysis buffer (methanol/acetonitrile/uLC-MS H_2_O (2:2:1)) was added per 1×10^6^ cells. Samples were vortexed and centrifuged at 14.000 x g for 15 minutes at 4°C. Supernatants were collected for LC-MS analysis.

### Liquid chromatography-mass spectrometry (LC-MS)-based metabolomics

All samples were analyzed by LC-MS using a Thermo Scientific™ Vanquish™ Horizon UHPLC System coupled to an Orbitrap Excedion Pro Mass Spectrometer. Metabolites were separated on a Sequant ZIC-pHILIC column (2.1 × 150 mm, 5 μm, Merck, Darmstadt, Germany) with a guard column (2.1 × 20 mm, 5 μm, Merck, Darmstadt, Germany) using a linear gradient with elution buffers of acetonitrile and 20 mM (NH4)2CO3, 0.1% NH4OH in ULC/MS grade water. Gradient ran from 20% eluent B to 60% eluent B in 20 minutes, followed by a wash step at 80% and equilibration at 20%, with a flow rate of 100 μL/min.

Mass spectrometric data were acquired in profile mode over an m/z range of 75-1000. Source settings were as follows: spray voltage, +3.5 kV in positive-ion mode and −2.5 kV in negative-ion mode; sheath gas, 35 arbitrary units; auxiliary gas, 10 arbitrary units; sweep gas, 0 arbitrary units; ion transfer tube temperature, 280 °C; vaporizer temperature, 150 °C; and RF lens, 50%. Full-scan MS^1^ data were acquired at a resolving power of 120,000 using a standard AGC target and automatic maximum injection time. For MS^1^-based metabolomics analyses, data were acquired using polarity switching. In either the custom or equidistant eDR workflows, the selected MS^1^ m/z range was subdivided into adjacent eDR windows, that overlap by 5 m/z within high-transmission regions of the quadrupole, ensuring that neighboring windows do not introduce an m/z gap or low-transmission boundary in the final spectrum. During spectral stitching, Advanced Peak Determination is utilized to improve determination of the charge states and monoisotopic m/z values of isotopic envelopes. If an isotope distribution is fully contained in only one window, the corresponding signals in the final spectrum are taken from that window. If an isotope distribution is fully contained in both overlapping windows, the signal intensities from both windows are compared, and the stitching boundary is placed within the overlap according to the window that provides the higher-intensity representation of that distribution. In this way, the cut-off decision is made inside the shared high-transmission overlap, preserving precursor isotope envelopes across window borders rather than truncating them at the nominal window edge. The independently acquired eDR subscans are then automatically stitched into a single full MS^1^ spectrum in the .raw file, preserving spectral quality while maintaining compatibility with downstream data processing and established workflows.

For eDR-enabled methods, two windowing strategies were assessed: an automatic mode using 12, 14 and 16 equidistant windows across the scanned m/z range and a user-defined mode with manually specified number of windows and customized window sizes (Supplemental Table 1).

For AcquireX intelligent data acquisition-based MS^2^ analyses, samples were acquired in separate positive- and negative-ion runs. Survey MS^1^ scans were acquired from m/z 75-1000 at 120,000 resolution, and MS^2^ spectra were acquired in data-dependent mode using AcquireX intelligent data acquisition-generated inclusion lists. Five ddMS^2^ scans were acquired per cycle using a 1.0 m/z isolation window and HCD with stepped normalized collision energies of 10, 35, and 60. MS^2^ spectra were acquired at a resolving power of 30,000 using a standard AGC target.

### Data analysis

Raw data were processed in Compound Discoverer (Thermo Fisher Scientific) versions 3.4/3.5 Software. Separate untargeted metabolomics workflows were used for MS^1^-only and AcquireX ddMS^2^ datasets. In both workflows, spectra were selected, retention times aligned, features detected and grouped, and compounds assembled prior to annotation and peak area assignment. Elemental composition prediction and ChemSpider searching were used for formula- and exact-mass-based annotation, and background features were filtered using the Mark Background node. For MS^2^ datasets, the workflow additionally incorporated mzCloud Mass Spectral Library matching and mzLogic-based prioritization of candidate annotations. In parallel, data were processed using TraceFinder 5.3 Software (Thermo Fisher Scientific), where metabolites were identified by matching to an in-house developed library, applying a mass tolerance of 5 ppm alongside retention time matching of standards.

Statistical analyses were performed using the R 4.4.2 and RStudio software. In all figure legends it is indicated which statistical tests were performed, as well as the p-value and the amount of technical and biological replicates.

## Results

### Full Scan eDR MS improves feature detection as compared to Full Scan MS acquisition

To systematically evaluate the impact of enhanced dynamic range (eDR) scanning mode on untargeted metabolomics performance, we analyzed lysates from the human HCC HepG2 cell line, comparing conventional Full Scan MS acquisition with Full Scan eDR MS acquisition mode with 12, 14 or 16 equidistant windows. Data files were processed in Compound Discoverer Software for feature detection and alignment across samples. Features with intensities at least threefold higher than blanks and a coefficient of variation (CV) < 20% across replicates were included in downstream analysis. Enabling eDR substantially increased the number of compound features compared to Full Scan MS acquisition, up to a 3.5-fold increase in detected features (Fig. 2A). A 12-window eDR configuration yielded the highest number of compound features (6965), followed by 16-window (6076) and 14 window (5726) setups, whereas conventional Full Scan MS mode resulted in the lowest number of compound features (1925) (Fig. 2A). Rank-intensity distributions demonstrated that all eDR conditions extended metabolite detection towards lower-abundance features, resulting in increased metabolite coverage (Fig. 2B, S1A). Notably, this effect was most pronounced for the 12-window configuration.

**Figure 2.**
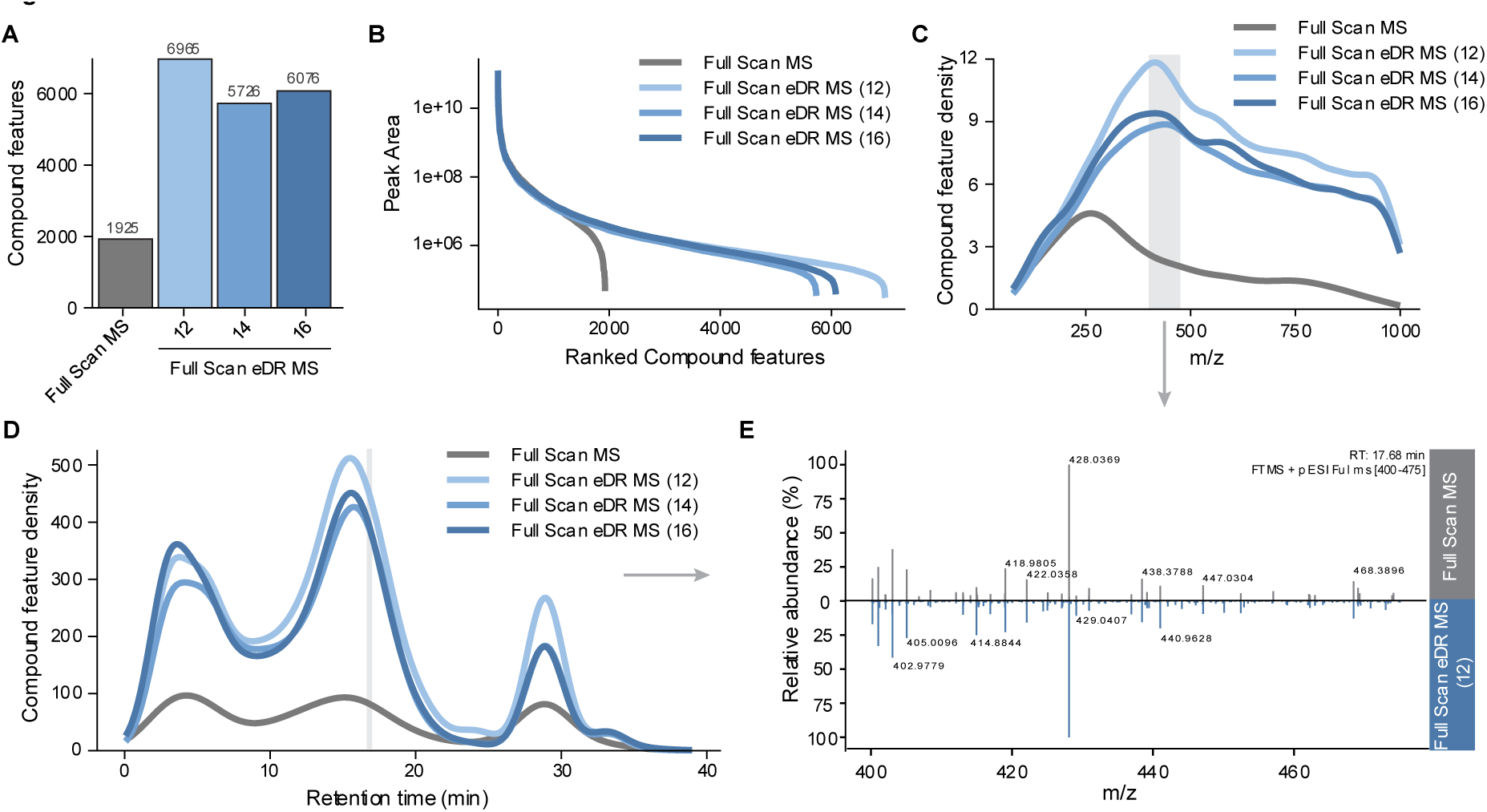
eDR scanning mode improves feature detection as compared to full-scan acquisition. Grouped compound features were detected in intracellular extracts of human HepG2 cells using Compound Discoverer Software for conventional Full Scan MS and Full Scan eDR MS, using 12, 14, or 16 equidistant isolation windows. Reported feature counts were filtered to retain only groups with signal intensities of at least threefold higher than the blanks and CV values below 20%. **A)** Total number of compound features for each acquisition mode. **B)** Rank-Intensity curve of compound features. **C)** Distribution of detected compound features across the m/z range. **D)** Distribution of detected compound features across chromatographic retention time (RT). **E)** A representative MS^1^ spectrum from the region with the highest feature density (m/z 400-475) at RT 17.5.

To determine where across the mass spectrum and chromatography the extra coverage of compound features in eDR occurs, we examined the distribution of detected compound features across conditions. The equidistant eDR acquisition modes predominantly increased detection in the higher m/z range (m/z > 500) (Fig. 2C). In addition, eDR enhanced feature detection in chromatographically dense regions (Fig. 2D), indicating improved coverage of co-eluting species as expected. To illustrate the spectral differences underlying eDR performance, a representative MS spectrum was extracted from the m/z region with the highest feature density (m/z 400-475, Fig. 2C) at the retention time around the maximal feature detection (17.7 min, Fig. 2D). Compared to Full Scan MS acquisition, Full Scan eDR MS enabled detection of numerous additional low-intensity ions that were either absent or suppressed in Full Scan MS, resulting in increased spectral depth and a more uniform signal distribution (Fig. 2E). Together, these results demonstrate that eDR acquisition increases analytical depth by revealing additional low-abundance features while preserving detection of abundant species, with the 12-window configuration providing the best overall performance for HILIC-based chromatography-MS of HepG2 cell lysates. Improvements were particularly detectable in the higher m/z range and in chromatographically dense regions.

### Customizing the amount and size of eDR windows allows for improved feature detection and sensitivity in predetermined m/z ranges

As many metabolites relevant to cancer biology fall into the lower m/z range (m/z <400), we next investigated whether eDR window design could be tuned to preferentially enhance detection in low mass regions. To this end, we designed five customized eDR acquisition schemes containing 12, 14 or 16 windows with non-uniform mass range distributions (Fig. 3A, Supplemental Table 1). In contrast to equidistant mass range window distributions, these customized eDR designs allocated narrower windows to the lower m/z region, where ion density was greatest. By portioning highly populated m/z regions into smaller segments, high-abundance ions were more effectively separated from lower-abundance species. This reduces the likelihood that ion accumulation is dominated by a small number of abundant ions, thereby enabling longer injection times and improved detection of low-abundance ions.

**Figure 3.**
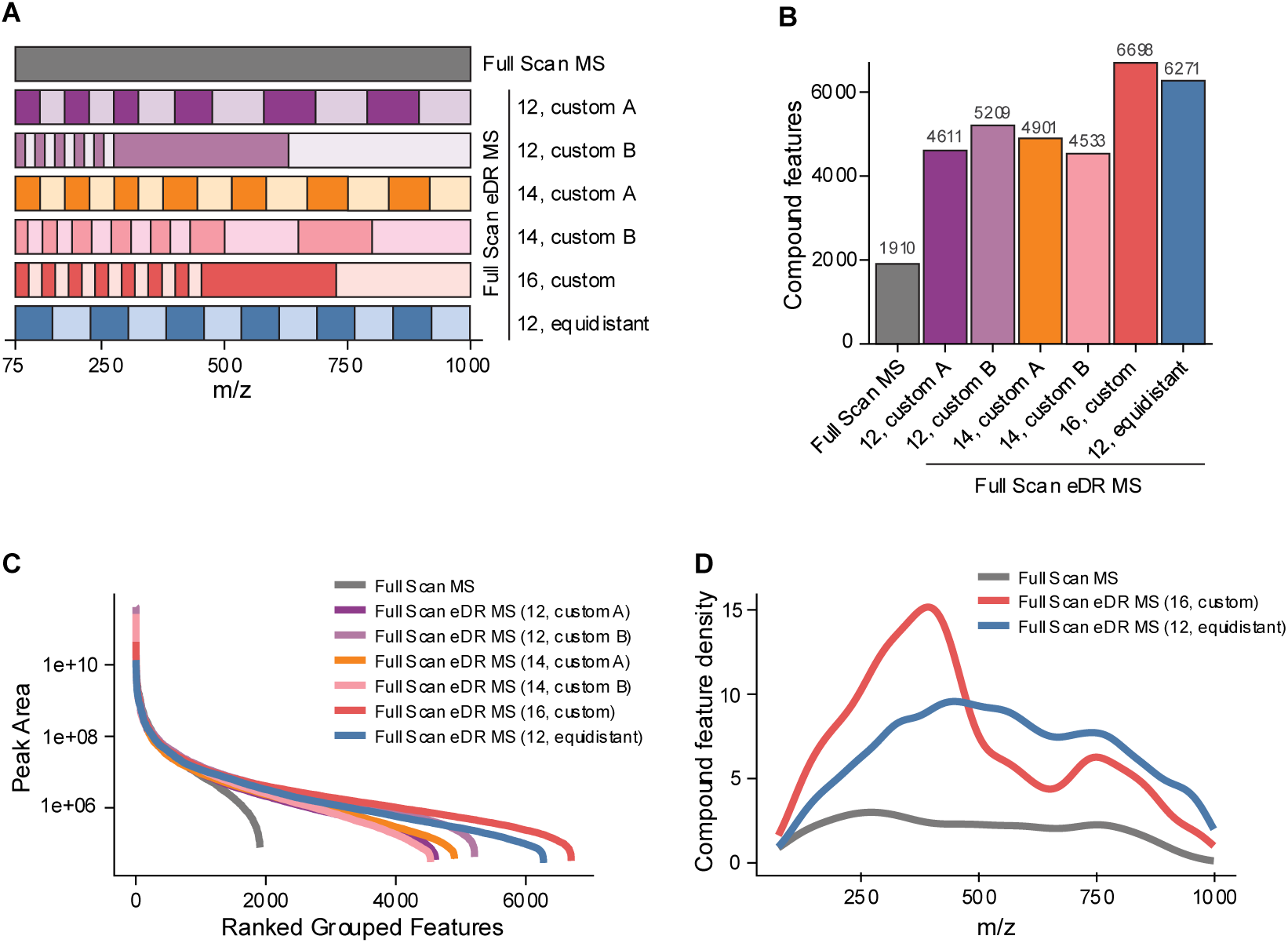
Customizing of eDR isolation windows improves feature detection within selected m/z ranges. **A)** Schematic overview of the customized eDR scan mode settings with 12, 14 or 16 windows with non-uniform isolation windows, including the eDR scanning mode with 12 equidistant windows and Full Scan MS. **B)** Total number of detected compound features in HepG2 extracts analyzed using different acquisition modes. Features were filtered to retain only those with signal intensities at least threefold higher than blank samples and CV values below 20% across 3 replicates. **C)** Rank-Intensity curve of compound features. **D)** Distribution of detected compound features across the m/z range.

For direct comparison, we also included a 12-window scanning mode with an equidistant distribution and a standard Full Scan MS mode. Using these acquisition modes, HepG2 lysates were analyzed and processed for feature detection and alignment. Consistent with earlier findings, all eDR enabled methods increased the total number of detected compound features relative to the Full Scan MS (2.4 – 3.5 fold change). Among all configurations, a customized 16-window design yielded the highest compound feature count (6698), followed by a 12-window equidistant configuration (6271) (Fig. 3B). All eDR conditions increased feature detection by extending detection towards lower-intensity features, with the customized 16-window and equidistant 12-window configurations exhibiting the broadest feature intensity distributions (Fig. 3C, S1B).

To determine whether customized window allocation selectively enhanced low-mass range coverage, compound features were ranked by m/z and compared across acquisition modes. Notably, customized eDR schemes with smaller window sizes allocated to the lower m/z region selectively enhanced the detection of low-mass features, illustrated by a steeper curve at the low m/z region (Fig. S1C). The m/z distribution corroborated this result, revealing a clear leftward shift in feature accumulation for the customized 16-window design (Fig. 3D). In contrast, the 12-window equidistant configuration, while showing high total feature count, did not specifically increase feature detection in the lower m/z range, but primarily extended detection in the higher m/z range. This confirms that targeted narrowing of eDR windows effectively redistributes acquisition sensitivity toward predefined mass ranges. Allocating narrower windows to the lower m/z region reduces dominance of high-abundance ions during ion accumulation in these segments, thereby improving detection of co-eluting low-abundance species. The 16-window customized configuration appears to achieve optimal balance between regional ion statistics and duty cycle, enabling both maximal total feature recovery and selective enhancement in the low-mass domain. These results demonstrate that eDR performance can be specifically tailored through window design. While equidistant segmentation maximizes global compound feature counts, customized window allocations enable targeted enhancement of metabolite detection within biologically relevant m/z regions.

### Customized eDR scanning mode improves feature detection and data quality in the lower m/z range

To assess the robustness and reproducibility of the optimized eDR strategy, we next evaluated performance across four hepatocellular carcinoma (HCC) cell lines: HepG2, Hep3B, Huh-7 and SNU-449, comparing a 16-window customized scheme to a 16-window equidistant eDR method and to conventional Full Scan MS. Across all cell lines, both eDR-enabled methods substantially increased the total number of detected features relative to Full Scan MS. The highest overall count was observed for the 16-window equidistant configuration, reaching up to 7249 features in SNU-449 (Fig. 4A). Consistently, both Full Scan eDR MS methods increased metabolite coverage by extending detection towards lower-intensity features across all cell lines, with the equidistant 16-window showing the broadest intensity distribution across all four cell lines (Fig. 4B, S2A).

**Figure 4.**
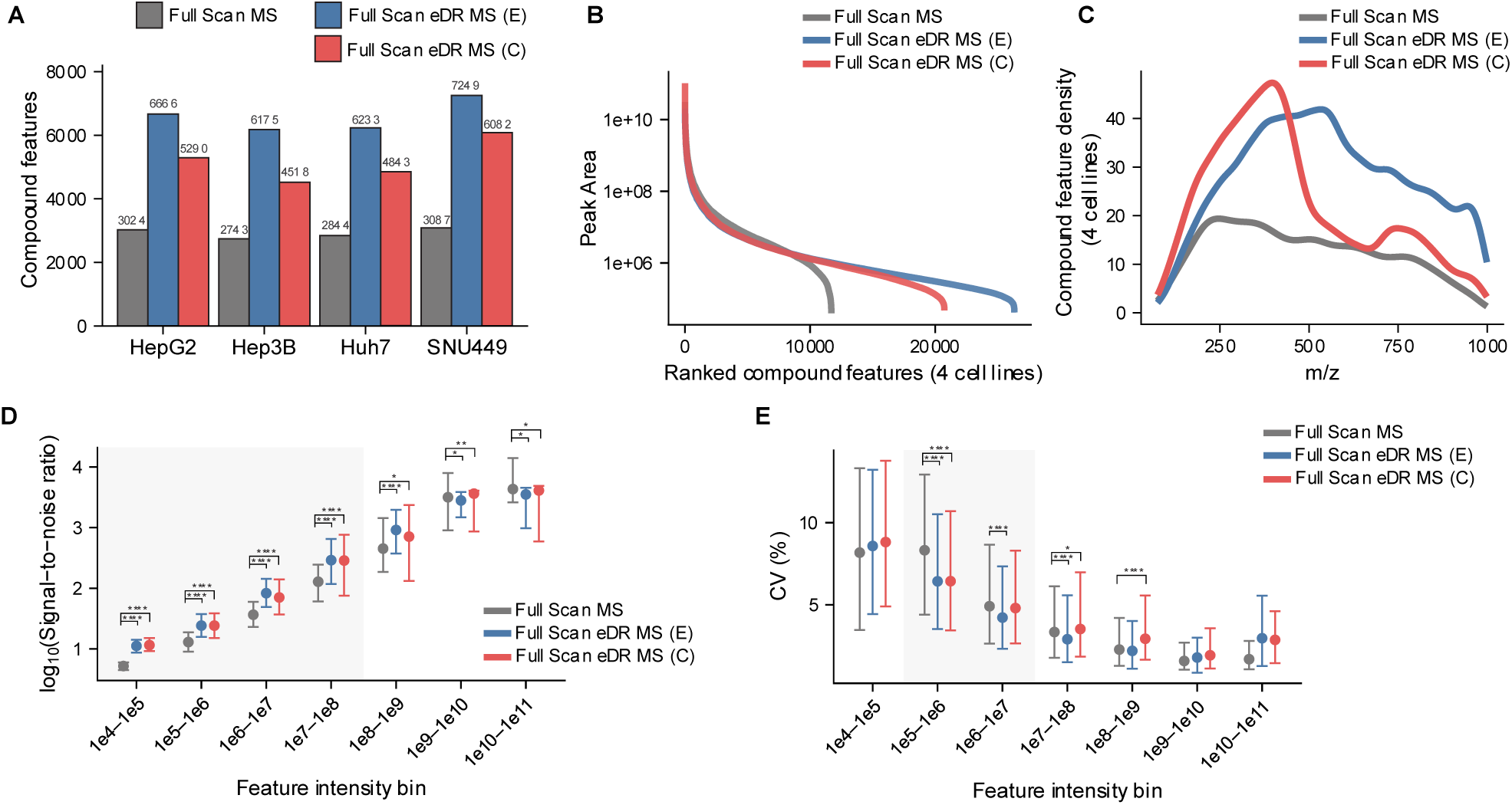
Customized eDR scanning mode improves feature detection and data quality in the lower m/z range. Four human hepatocellular carcinoma (HCC) intracellular matrices (HepG2, Hep3B, Huh-7, SNU-449) were analyzed using Full Scan MS and Full Scan eDR MS, using 16 windows with either a customized version (C) or with an equidistant distribution (E) (n=3). **A)** Total number of detected compound features for all 4 cell lines. **B)** Rank-Intensity curve of compound features for all 4 cell lines combined. **C)** Distribution of detected compound features across the m/z range for all 4 cell lines combined. **D)** Signal-to-noise ratios (S/N, log10-transformed) of detected compound features across intensity bins for the three acquisition modes. Data from all four HCC cell lines were combined. **E)** Coefficients of variation (CV) of detected compound features across intensity bins for the three acquisition modes. Data from all four HCC cell lines were combined. Within each intensity bin and cell line, pairwise differences between scan modes were assessed using two-sided Wilcoxon rank-sum tests. P-values were adjusted for multiple comparisons using the Benjamini–Hochberg method (* = p<0.05, **= p<0.01, ***= p<0.001, ****= p<0.0001).

We next examined whether the customized windows design preserved its selective enhancement of low m/z detection across cell lines. Consistent with our earlier findings, customized eDR scanning mode showed a steeper accumulation in feature detection in the lower m/z range compared to both equidistant Full Scan eDR MS and Full Scan MS modes (Fig. 4C, Fig. S2C). Visualization of m/z distributions confirmed a consistent enrichment of low-mass features for the customized acquisition strategy across cell lines (Fig. 4C, S2B). While equidistant Full Scan eDR MS showed the strongest increase in compound feature detection, its strongest gains were observed in the higher m/z region (Fig. 4C). In contrast, the customized eDR scanning mode specifically performed well in the low m/z range (Fig. 4C). Importantly, these patterns were conserved across all four individual cell lines (Fig. S2B, S2C), demonstrating that mass-selective enhancement via window customization is matrix-independent.

Beyond feature detection, we evaluated several data quality metrics to determine whether eDR acquisition improves qualitative performance. Features were stratified into intensity bins, and signal-to-noise ratios (S/N) and coefficients of variation (CVs) were calculated from triplicate measurements. Signal-to-noise analysis revealed significant increases in S/N for low-intensity features (< 1e8) using eDR acquisition (Fig. 4D, S2D). In addition, both eDR methods significantly reduced CV values relative to Full Scan MS in the 1e5-1e6 and 1e6-1e7 intensity bins (Fig. 4E, S2E). While the lowest intensity bin (1e4-1e5) showed higher S/N in Full Scan eDR MS versus Full Scan MS modes, there was no reduction in CV, probably due to the low number of features in this bin and the high variation in low-abundance species. High intensity features exhibited no significant changes in CV values, as CVs were already low under Full Scan MS conditions (<5%) (Fig. 4E). Collectively, these findings demonstrate that Full Scan eDR MS acquisition significantly increases feature detection and metabolite coverage across independent cell lines and that customization of the eDR windows can enhance certain m/z regions of interest, all while not compromising the quantitative performance of the instrument.

### (Customized) eDR scanning mode improves MS^2^ coverage and identification of cancer-relevant metabolites

We next evaluated whether the improved MS^1^ performance of Full Scan eDR MS translated into enhanced MS^2^ acquisition and metabolite identification. To this end, a mixed pool of cell lysates was subjected to iterative AcquireX intelligent data acquisition. When using standard Full Scan MS acquisition in the data-dependent AcquireX deep scan workflow, detectable precursors were rapidly exhausted, causing the number of newly acquired MS^2^ spectra to plateau early in both positive and negative mode (Fig. S3A). In contrast, Full Scan eDR MS acquisition markedly delayed this exhaustion, enabling sustained fragmentation of new precursors across repeated injections, indicative of deeper fragmentation coverage under eDR conditions. Indeed, both eDR-enabled methods increased the number of MS^2^ spectra across cell lines (Fig. 5A).

**Figure 5.**
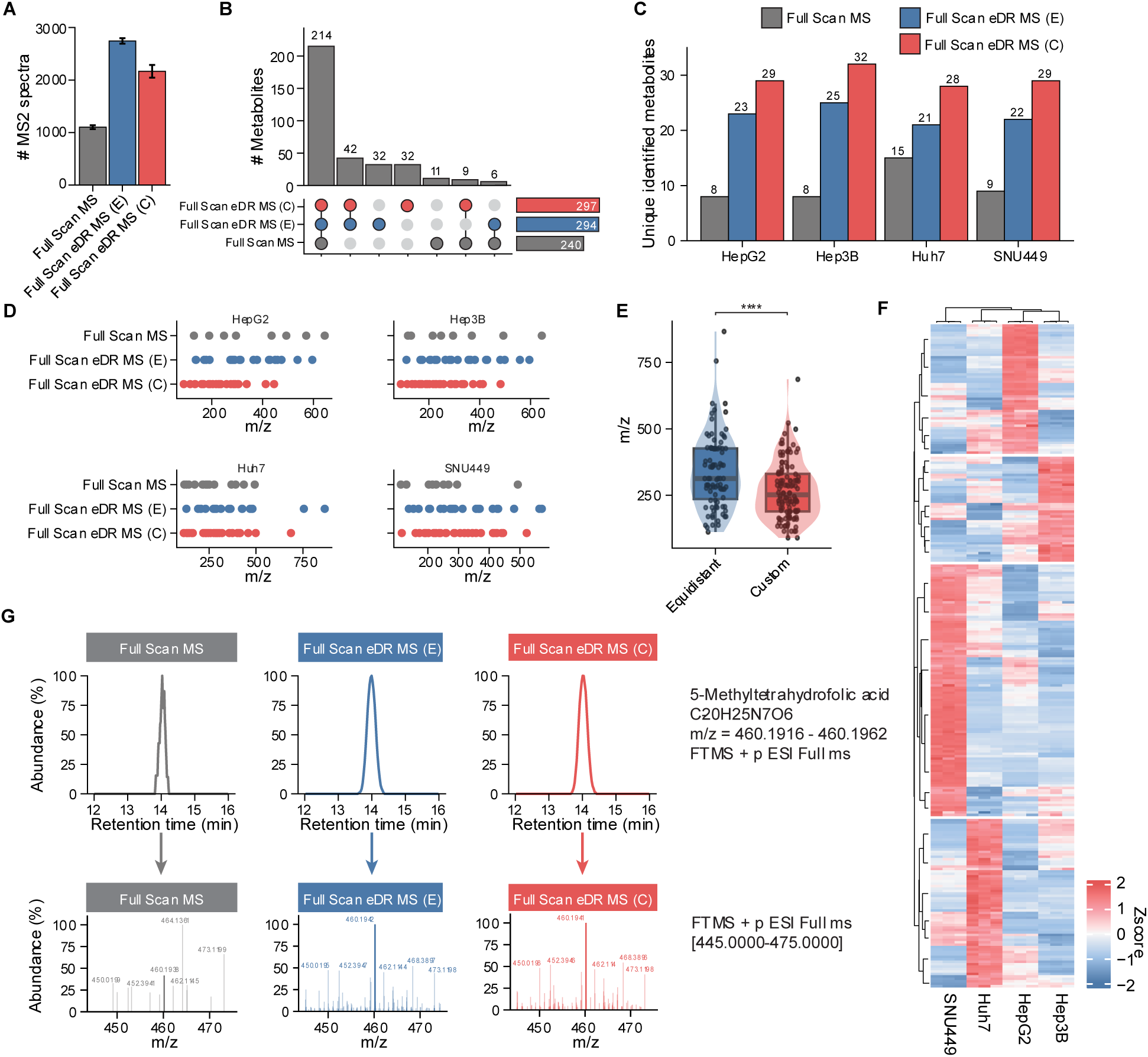
(Customized) eDR acquisition improves MS^2^ coverage and identification of cancer-relevant metabolites. MS^2^ acquisition and metabolite identification was analyzed in a pooled sample of the four human HCC intracellular matrices (HepG2, Hep3B, Huh-7, SNU-449), using the data-dependent AcquireX deep scan workflow with Full Scan MS and Full Scan eDR MS, using 16 windows with either a customized version (C) or with an equidistant distribution (E). **A)** Total number of acquired MS^2^ spectra for each acquisition method across all HCC cell lines. **B)** UpSet plot showing metabolite identifications for each acquisition method across all HCC cell lines. Metabolite annotations were generated using mzCloud-based matching in Compound Discoverer Software with a match factor threshold of 60, and targeted analysis using TraceFinder and an in-house library (n=140). Metabolites present in 3 out of 3 replicates per cell line and CV values below 20% were included for downstream analysis. Horizontal bars represent the total number of identified metabolites per acquisition method, whereas vertical bars indicate the number of metabolites uniquely or jointly identified per acquisition mode. **C)** Number of uniquely identified metabolites per acquisition method for each individual cell line. **D)** Distribution of m/z values for uniquely identified metabolites across acquisition methods and cell lines. Each dot represents a single metabolite. **E)** A violin-box plot representing the distribution of m/z values for uniquely identified metabolites in the customized and equidistant-distributed 16-window eDR scanning modes across all four HCC cell lines. The box represents the interquartile range (IQR) with the central horizontal line indicating the median, and whiskers extend to 1.5x IQR. Statistical difference between groups was assessed using a two-sided Wilcoxon rank-sum test (p=6e^-5^). **F)** Extracted ion chromatogram (XIC) of the low-abundance one-carbon metabolite 5-methyltetrahydrofolic acid (5-MTHF) (top) and corresponding MS spectra (bottom). **G)** Hierarchically clustered heatmap of significantly altered metabolites across HCC cell lines is displayed based on z-scores-transformed metabolite abundances.

To assess metabolite identification performance, we compiled a high-confidence identification dataset using complementary approaches: (i) mzCloud Mass Spectral Library-based annotations generated in Compound Discoverer Software with a match factor threshold of 60, (ii) manual curation of these annotations, (iii) targeted annotation using an in-house library (n=140). These datasets were combined into a final curated identification list for downstream comparisons, filtered on presence in 3 out of 3 replicates per cell line and CV < 20%. Using this curated dataset, we observed a substantial increase in confident metabolite identification with Full Scan eDR MS acquisition compared to Full Scan MS. With 297 identified metabolites across all cell lines, the customized eDR mode achieved the highest identified metabolite count, while Full Scan MS enabled 240 metabolite identifications (Fig. 5B). The same trend was present at the individual cell line level, with the customized eDR mode consistently resulting in a higher number of metabolite identifications in all cell lines (Fig. S3B).

While 214 metabolites were identified in all three scan modes across cell lines, both eDR modes resulted in 32 uniquely identified metabolites and share 42 metabolite identifications that were not present in Full Scan MS (Fig. 5B). Additionally, the customized eDR mode consistently showed higher unique identifications as compared to the equidistant eDR mode in the individual cell lines (Fig. 5C), indicating that the customized eDR mode is able to identify most unique metabolites in four cell lines. Importantly, many uniquely identified metabolites under customized eDR acquisition resided in the lower m/z range in all cell lines (Fig. 5D), resulting in significantly lower m/z values of uniquely identified metabolites in eDR custom as compared to the equidistant eDR mode (Fig. 5E). These data underscore the added value of customized windows to enhance detection and identification in the low m/z range.

Full Scan MS resulted in only 11 uniquely identified metabolites (Fig. 5B). Detailed inspection revealed that only two of these identifications represented clear Full Scan MS-specific detections. In these cases, the affected metabolites were low-abundance ions co-isolated with high-abundance species within the corresponding eDR windows. In these situations, a reduced injection time for the mass range will not only reduce S/N of the high-abundance compounds but all compounds which elute within this RT and m/z window. For the remaining cases, precursor signals were also detected in the eDR datasets, but were not converted to final metabolite identifications due to differences in MS^2^ acquisition, compound annotation or downstream data processing. These findings indicate that the small number of metabolites uniquely identified by Full Scan MS reflects isolated acquisition- or processing-dependent effects rather than an inherent limitation of eDR. Importantly, the limited number of metabolites preferentially identified using Full Scan MS had little impact on the overall biological interpretation of the dataset when compared with the substantial increase in metabolite identifications achieved using eDR.

Beyond individual metabolite detection, the increased analytical depth achieved with eDR translated into improved resolution of metabolic differences between cell lines. The customized Full Scan eDR MS method yielded 268 significantly altered annotated metabolites across the HCC cell lines, compared with 258 for the equidistant Full Scan eDR MS approach and 215 for Full Scan MS (adjusted p-value<0.05) (Fig. S3C). Unsupervised clustering based on these metabolites revealed clear stratification of the different cell lines, reflecting their underlying metabolic heterogeneity (Fig. 5F).

Notably, 71 metabolites were only identified as significantly altered using the customized Full Scan eDR MS method compared to Full Scan MS. Only 12 of these metabolites were also annotated in Full Scan MS, indicating that the enhanced biological resolution provided by eDR largely originated from the identification of metabolites that could not be confidently annotated using conventional Full Scan MS. These metabolites included compounds involved in biological relevant pathways associated with cancer metabolism, including one-carbon metabolism, nucleotide metabolism, and the mevalonate pathway.

Among these were low-abundance 1C metabolites, including folate derivatives such as 5-methyltetrahydrofolate (5-MTHF), that was only consistently detected under eDR conditions (Fig. 5G), highlighting the ability of eDR to capture metabolites that are typically near or below detection limits in complex matrices. In addition, key intermediates of the mevalonate pathway, including mevalonate and HMG-CoA were only detected using eDR. This pathway is critical for cholesterol biosynthesis and protein prenylation, processes that are frequently upregulated in cancer and have been specifically implicated in HCC progression and proliferation. Given the important roles of these pathways in cancer cell proliferation and HCC progression, these findings demonstrate that the improved analytical performance of eDR translates directly into improved biological insight.

Together, these results show that eDR acquisition substantially increased MS^2^ coverage and metabolite identifications in an untargeted metabolomics approach. In addition, eDR enables quantification of biologically important metabolic pathways such as 1C metabolism and the mevalonate pathway and improves the resolution of cancer-associated metabolic heterogeneity. The ability to customize window size and distribution provides a flexible strategy to enhance the detection of low-abundance metabolites in predefined mass regions, supporting deeper and targeted interrogation of complex biological systems.

## Discussion

LC-MS based untargeted metabolomics has proven powerful for metabolite profiling and the discovery of novel metabolites across diverse biological samples. However, untargeted metabolomics is limited by the wide concentration range of biological metabolites, which results in reduced sensitivity and metabolite coverage of low-abundance ions, obscuring biologically relevant signals (Cui et al., 2018; David et al., 2021; Meister et al., 2021).

Here, we demonstrate that Full Scan eDR MS acquisition substantially improves detection, quantification and identification of low-abundance metabolites in biological samples as compared to conventional Full Scan MS. By segmentation of the m/z domain, eDR increases effective ion accumulation for low-abundance species, thereby improving ion statistics and measurement precision in the low-intensity signals.

In conventional Full Scan MS, high-abundance ions dominate space-charge capacity in the C-trap, leading to rapid saturation and limiting the effective injection time available for low-abundance ions (Makarov, 2026; Makarov & Scigelova, 2010). As a result, low-intensity ions are disproportionately affected by ion suppression, leading to reduced detectability and reproducibility. eDR redistributes ion accumulation across the mass range by segmenting the full m/z range into two multiplexed Orbitrap subscans comprising customizable m/z windows. Within each subscan, multiple m/z windows are non-sequentially isolated and accumulated into a single orbitrap injection. By reducing the contribution of high-abundance ions within individual m/z segments, eDR is expected to improve the detectability of lower-abundance species while maintaining detection of abundant metabolites.

Our data show that overall, eDR acquisition has several important downstream benefits. First, the increased ion trapping for low-abundance features resulted in improved S/N ratios, leading to a higher number of detected metabolites and a deeper profiling of the metabolome in complex samples. In addition, the increased signal quality improved precursor ion selection for MS^2^, enhancing metabolite identification. As a result, eDR provides a more in-depth analysis of the metabolome without requiring additional sample preparation or fractionation strategies. By adjusting the number, size and distribution of m/z windows, the acquisition scheme can be further tailored to the ion density profile and region of interest of a given sample. In our samples, this proved particularly advantageous in chromatographically dense regions, where co-eluting compounds suffer from C-trap capacity limitations. Allocating narrow windows to the low m/z region, which was most densely populated in our samples, allowed better control over ion injection times and improved detection of low mass ions. In general, optimal performance of Full Scan eDR MS depends on the chromatography and sample complexity. Window design should be adapted to the specific analytical context and metabolites of interest to achieve the best results. In addition, segmentation of the m/z domain introduces potential slower scan speed and higher duty cycle. Optimization is therefore required to balance dynamic range extension with resolution and chromatographic peak width. Importantly, we show that the gains in analytical performance of eDR translate into improved metabolite coverage of biologically relevant pathways. Low-abundance intermediates from 1C metabolism, including 5-MTHF, are frequently underrepresented in conventional LC-MS workflows due to their low abundance, but were consistently detected under eDR conditions. These metabolites play a central role in folate-mediated carbon transfer reactions that support nucleotide biosynthesis, methylation capacity and redox homeostasis and are linked to cancer progression. Importantly, alterations in 1C metabolism have been associated with HCC progression, primarily evidenced by dysregulated enzymes within this pathway. Overexpression of 1C enzymes is linked to poor patient prognosis, while inhibition of 1C resulted in decreased cell growth rates, underscoring its clinical relevance and therapeutic potential in HCC (Guerrero et al., 2022; Quevedo-Ocampo et al., 2022; Ren et al., 2022). The current understanding is however largely limited to enzyme-level alterations, as 1C metabolites remain insufficiently characterized to date. This represents a previously unexplored layer that could provide valuable insights into 1C metabolic regulation, which may reveal potential novel targets in HCC. Metabolites related to the mevalonate pathway, including mevalonate and HMG-CoA, also showed improved detectability. This pathway is essential for cholesterol biosynthesis, protein prenylation and CoQ synthesis and has been widely linked to cancer cell proliferation and survival, including of HCC (Chen et al., 2025; Liang et al., 2019). The ability to analyze these metabolites highlights that eDR does not merely increase the number of detected features but can also enable interrogation of biologically meaningful metabolic layers that are typically obscured using conventional acquisition strategies. While the current study does not aim to provide a detailed interpretation of these differences, our data underscore the potential of Full Scan eDR MS to improve metabolomics analysis of biologically relevant, low-abundance metabolites in heterogenous systems. This likely becomes increasingly important when analyzing complex matrices, such as tissues, biofluids or microbiome-derived samples, which are generally characterized by greater chemical diversity, broader dynamic range and more unknown compounds as compared to cell-line samples. While future studies are needed to evaluate the performance of eDR in such complex samples, we expect that due to its increased sensitivity and ability to identify more metabolites, eDR will show equal or higher benefits in such complex matrices.

Full Scan eDR MS complements and extends existing strategies aimed at improving metabolite coverage and identification. Approaches such as BoxCar acquisition similarly segment the m/z domain to mitigate dynamic range limitation. Iterative inclusion/exclusion strategies, implemented in AcquireX intelligent data acquisition workflows, can further improve MS^2^ coverage by reducing redundant fragmentation of repeatedly selected precursor ions. However, these strategies remain fundamentally dependent on the completeness of the initial MS^1^ feature inventory. We here show that using conventional full-scan acquisition in AcquireX intelligent data acquisition workflows resulted in a rapid exhaustion of detectable precursors, causing the number of newly acquired MS^2^ spectra to plateau early. Likely, low-and medium-abundance compounds remained undetected or insufficiently represented at the MS^1^ level in this case and were therefore not transferred into subsequent inclusion/exclusion lists. By increasing MS^1^-level detectability across a wider dynamic range, eDR improves the visibility of these lower-abundance precursors. In AcquireX intelligent data acquisition workflows, inclusion lists specifically target these newly detected features for fragmentation, resulting in the acquisition of MS^2^ spectra for a larger and more representative population of features that would otherwise be missed in standard full-scan MS^1^ acquisition. Consequently, the benefit of eDR is not restricted to improved MS^1^ profiling but also translates into enhanced MS^2^ coverage through the acquisition of additional MS^2^ spectra for compounds that become detectable only under eDR conditions.

In conclusion, our work establishes Full Scan eDR MS as a versatile acquisition strategy for untargeted metabolomics. The compatibility of eDR with existing LC-MS workflows combined with a tunable design makes it broadly applicable across diverse experimental settings. By improving ion utilization efficiency at the MS^1^ level and enhancing downstream MS^2^ acquisition, eDR increases metabolite coverage, analytical depth and metabolite identification while preserving data quality. Importantly, these gains translate into improved detection of biologically relevant low-abundance metabolite that would otherwise remain inaccessible. As such, Full Scan eDR MS provides a practical framework for deeper interrogation of complex biological systems by expanding metabolome coverage and improving the detection of biologically relevant low-abundance metabolites.

## Supporting Information

Detailed Full Scan eDR MS acquisition window configurations and m/z boundaries for all evaluated scan modes (Table S1); supplementary analyses of metabolite feature detection across eDR acquisition modes, metabolite m/z distributions and supplementary performance comparisons. (Figures S1-S3) (PDF).

## Acknowledgements

The authors would like to thank Claire Dauly (Thermo Fisher Scientific, Bremen, Germany) for valuable advice and constructive contributions to the study design.

## Author Contributions

DR, MK, KF, SB, CB and EZ conceived the project and designed the experiments. DR, MK and CK performed experiments and LC-MS analysis. CT developed the eDR acquisition mode used in this study. DR and EZ conducted data analysis and interpretation. SB, CB and EZ supervised the study. DR, MK, SB, CB, EZ, wrote the manuscript, which was reviewed by all co-authors. All authors have given approval to the final version of the manuscript.

## Declaration of interests

Susan Bird, Michał Kaczmarek, Christian Klaas, Kyle Fort and Christian Thoeing are employees of Thermo Fisher Scientific (manufacturer of Orbitrap mass spectrometers). All other authors declare no conflicts of interest.

## References

Alseekh, S., Aharoni, A., Brotman, Y., Contrepois, K., D’Auria, J., Ewald, J., C. Ewald, J., Fraser, P. D., Giavalisco, P., Hall, R. D., Heinemann, M., Link, H., Luo, J., Neumann, S., Nielsen, J., Perez de Souza, L., Saito, K., Sauer, U., Schroeder, F. C., … Fernie, A. R. (2021). Mass spectrometry-based metabolomics: A guide for annotation, quantification and best reporting practices. Nature Methods, 18(7), 747–756. 10.1038/s41592-021-01197-1

Cai, M., Liu, H., Shao, C., Li, T., Jin, J., Liang, Y., Wang, J., Cao, J., Yang, B., He, Q., Shao, X., & Ying, M. (2025). Metabolomics and metabolites in cancer diagnosis and treatment. Molecular Biomedicine, 6, 109. 10.1186/s43556-025-00362-8

Chen, Y., Lee, D., Kwan, K. K.-L., Wu, M., Wang, G., Zhang, M. S., Deng, H., Cheu, J. W.-S., Lau, M. H.-Y., Chan, C. Y.-K., Ooi, Z. Y., Wu, Y., Bao, M. H.-R., Lo, R. C.-L., Ng, I. O.-L., Wong, C.-M., & Wong, C. C.-L. (2025). Mevalonate pathway promotes liver cancer by suppressing ferroptosis through CoQ10 production and selenocysteine-tRNA modification. Journal of Hepatology, 83(6), 1338–1352. 10.1016/j.jhep.2025.06.034

Cooper, B., & Yang, R. (2024). An assessment of AcquireX and Compound Discoverer software 3.3 for non-targeted metabolomics. Scientific Reports, 14(1), 4841. 10.1038/s41598-024-55356-3

Cui, L., Lu, H., & Lee, Y. H. (2018). Challenges and emergent solutions for LC-MS/MS based untargeted metabolomics in diseases. Mass Spec Rev. 2018;37:772–792. 10.1002/mas.21562

David, L., Kang, J., & Chen, S. (2021). Untargeted Metabolomics of Arabidopsis Stomatal Immunity. In J. J. Sanchez-Serrano & J. Salinas (Eds.), Arabidopsis Protocols (pp. 413–424). Springer US. 10.1007/978-1-0716-0880-7_20

Du, D., Liu, C., Qin, M., Zhang, X., Xi, T., Yuan, S., Hao, H., & Xiong, J. (2022). Metabolic dysregulation and emerging therapeutical targets for hepatocellular carcinoma. Acta Pharmaceutica Sinica. B, 12(2), 558–580. 10.1016/j.apsb.2021.09.019

Fernie, A. R., Trethewey, R. N., Krotzky, A. J., & Willmitzer, L. (2004). Metabolite profiling: From diagnostics to systems biology. Nature Reviews Molecular Cell Biology, 5(9), 763–769. 10.1038/nrm1451

Furlani, I. L., da Cruz Nunes, E., Canuto, G. A. B., Macedo, A. N., & Oliveira, R. V. (2021). Liquid Chromatography-Mass Spectrometry for Clinical Metabolomics: An Overview. In A. V. Colnaghi Simionato (Ed.), Separation Techniques Applied to Omics Sciences: From Principles to Relevant Applications (pp. 179–213). Springer International Publishing. 10.1007/978-3-030-77252-9_10

Gelambi, M., & Whitehead, S. R. (2024). Untargeted Metabolomics Reveals Fruit Secondary Metabolites Alter Bat Nutrient Absorption. Journal of Chemical Ecology, 50(7), 385–396. 10.1007/s10886-024-01503-z

González-Domínguez, Á., Armeni, M., Savolainen, O., Lechuga-Sancho, A. M., Landberg, R., & González-Domínguez, R. (2023). Untargeted Metabolomics Based on Liquid Chromatography–Mass Spectrometry for the Analysis of Plasma and Erythrocyte Samples in Childhood Obesity. In R. González-Domínguez (Ed.), Mass Spectrometry for Metabolomics (pp. 115–122). Springer US. 10.1007/978-1-0716-2699-3_11

Guerrero, L., Paradela, A., & Corrales, F. J. (2022). Targeted Proteomics for Monitoring One-Carbon Metabolism in Liver Diseases. Metabolites, 12(9). 10.3390/metabo12090779

Guijas, C., Montenegro-Burke, J. R., Warth, B., Spilker, M. E., & Siuzdak, G. (2018). Metabolomics activity screening for identifying metabolites that modulate phenotype. Nature Biotechnology, 36(4), 316–320. 10.1038/nbt.4101

Hanahan, D. (2026). Hallmarks of cancer—Then and now, and beyond. Cell, 189(8), 2254–2277. 10.1016/j.cell.2025.12.049

Huang, H., Chen, Y., Xu, W., Cao, L., Qian, K., Bischof, E., Kennedy, B. K., & Pu, J. (2025). Decoding aging clocks: New insights from metabolomics. Cell Metabolism, 37(1), 34–58. 10.1016/j.cmet.2024.11.007

Kaufmann, A., Maden, K., & Walker, S. (2020). Partially overlapping sequential window acquisition of all theoretical mass spectra: A methodology to improve the spectra quality of veterinary drugs present at low concentrations in highly complex biological matrices. Rapid Communications in Mass Spectrometry, 34(7), e8638. 10.1002/rcm.8638

Kennedy, A. D., Wittmann, B. M., Evans, A. M., Miller, L. A. D., Toal, D. R., Lonergan, S., Elsea, S. H., & Pappan, K. L. (2018). Metabolomics in the clinic: A review of the shared and unique features of untargeted metabolomics for clinical research and clinical testing. Journal of Mass Spectrometry, 53(11), 1143–1154. 10.1002/jms.4292

Kharazian, N., Dehkordi, F. J., & Xiang, C.-L. (2025). Metabolomics-based profiling of five Salvia L. (Lamiaceae) species using untargeted data analysis workflow. Phytochemical Analysis, 36(1), 113–143. 10.1002/pca.3423

Liang, B., Chen, R., Song, S., Wang, H., Sun, G., Yang, H., Jing, W., Zhou, X., Fu, Z., Huang, G., & Zhao, J. (2019). ASPP2 inhibits tumor growth by repressing the mevalonate pathway in hepatocellular carcinoma. Cell Death & Disease, 10(11), 830. 10.1038/s41419-019-2054-7

Liu, J., Geng, W., Sun, H., Liu, C., Huang, F., Cao, J., Xia, L., Zhao, H., Zhai, J., Li, Q., Zhang, X., Kuang, M., Shen, S., Xia, Q., Wong, V. W.-S., & Yu, J. (2022). Integrative metabolomic characterisation identifies altered portal vein serum metabolome contributing to human hepatocellular carcinoma. 10.1136/gutjnl-2021-325189

Lu, W., Su, X., Klein, M. S., Lewis, I. A., Fiehn, O., & Rabinowitz, J. D. (2017). Metabolite Measurement: Pitfalls to Avoid and Practices to Follow. Annual Review of Biochemistry, 86(Volume 86, 2017), 277–304. 10.1146/annurev-biochem-061516-044952

Makarov, A. (2026). First 20 Years of Orbitrap Mass Spectrometry as the Mainstream Analytical Technique. Mass Spectrometry Reviews. 10.1002/mas.70024

Makarov, A., & Scigelova, M. (2010). Coupling liquid chromatography to Orbitrap mass spectrometry. *Journal of Chromatography A*, Mass Spectrometry: Innovation and Application. Part VI, 1217(25), 3938–3945. 10.1016/j.chroma.2010.02.022

Meier, F., Geyer, P. E., Virreira Winter, S., Cox, J., & Mann, M. (2018). BoxCar acquisition method enables single-shot proteomics at a depth of 10,000 proteins in 100 minutes. Nature Methods, 15(6), 440–448. 10.1038/s41592-018-0003-5

Meister, I., Zhang, P., Sinha, A., Sköld, C. M., Wheelock, Å. M., Izumi, T., Chaleckis, R., & Wheelock, C. E. (2021). High-Precision Automated Workflow for Urinary Untargeted Metabolomic Epidemiology. Analytical Chemistry, 93(12), 5248–5258. 10.1021/acs.analchem.1c00203

Mishra, S., Maltseva, A., Nieminen, A. I., Niku, M., Karikka, S., Hekkala, J., Leppä, S., Vihinen, P., Sunela, K., Koivunen, J., Jukkola, A., Kalashnikov, I., Auvinen, P., Kääriäinen, O.-S., Saarnio, J., Meriläinen, S., Rautio, T., Aro, R., Häivälä, R., … Reunanen, J. (2025). The metabolome of fecal extracellular vesicles in patients with malignant solid tumors. Scientific Reports, 15, 29402. 10.1038/s41598-025-14250-2

Moreau, R., Clària, J., Aguilar, F., Fenaille, F., Lozano, J. J., Junot, C., Colsch, B., Caraceni, P., Trebicka, J., Pavesi, M., Alessandria, C., Nevens, F., Saliba, F., Welzel, T. M., Albillos, A., Gustot, T., Fernández, J., Moreno, C., Baldassarre, M., … Angeli, P. (2020). Blood metabolomics uncovers inflammation-associated mitochondrial dysfunction as a potential mechanism underlying ACLF☆. Journal of Hepatology, 72(4), 688–701. 10.1016/j.jhep.2019.11.009

Park, S., & Hall, M. N. (2025). Metabolic reprogramming in hepatocellular carcinoma: Mechanisms and therapeutic implications. Experimental & Molecular Medicine, 57(3), 515–523. 10.1038/s12276-025-01415-2

Pavlova, N. N., Zhu, J., & Thompson, C. B. (2022). The hallmarks of cancer metabolism: Still emerging. Cell Metabolism, 34(3), 355–377. 10.1016/j.cmet.2022.01.007

Pezzatti, J., Boccard, J., Codesido, S., Gagnebin, Y., Joshi, A., Picard, D., González-Ruiz, V., & Rudaz, S. (2020). Implementation of liquid chromatography–high resolution mass spectrometry methods for untargeted metabolomic analyses of biological samples: A tutorial. Analytica Chimica Acta, 1105, 28–44. 10.1016/j.aca.2019.12.062

Quevedo-Ocampo, J., Escobedo-Calvario, A., Souza-Arroyo, V., Miranda-Labra, R. U., Bucio-Ortiz, L., Gutiérrez-Ruiz, M. C., Chávez-Rodríguez, L., & Gomez-Quiroz, L. E. (2022). Folate Metabolism in Hepatocellular Carcinoma. What Do We Know So Far? Technology in Cancer Research & Treatment, 21, 15330338221144446. 10.1177/15330338221144446

Ranninger, C., Schmidt, L. E., Rurik, M., Limonciel, A., Jennings, P., Kohlbacher, O., & Huber, C. G. (2016). Improving global feature detectabilities through scan range splitting for untargeted metabolomics by high-performance liquid chromatography-Orbitrap mass spectrometry. Analytica Chimica Acta, 930, 13–22. 10.1016/j.aca.2016.05.017

Ren, X., Rong, Z., Liu, X., Gao, J., Xu, X., Zi, Y., Mu, Y., Guan, Y., Cao, Z., Zhang, Y., Zeng, Z., Fan, Q., Wang, X., Pei, Q., Wang, X., Xin, H., Li, Z., Nie, Y., Qiu, Z., … Deng, Y. (2022). The Protein Kinase Activity of NME7 Activates Wnt/β-Catenin Signaling to Promote One-Carbon Metabolism in Hepatocellular Carcinoma. Cancer Research, 82(1), 60–74. 10.1158/0008-5472.CAN-21-1020

Rinschen, M. M., Ivanisevic, J., Giera, M., & Siuzdak, G. (2019). Identification of bioactive metabolites using activity metabolomics. Nature Reviews Molecular Cell Biology, 20(6), 353–367. 10.1038/s41580-019-0108-4

Sarvin, B., Lagziel, S., Sarvin, N., Mukha, D., Kumar, P., Aizenshtein, E., & Shlomi, T. (2020). Fast and sensitive flow-injection mass spectrometry metabolomics by analyzing sample-specific ion distributions. Nature Communications, 11(1), 3186. 10.1038/s41467-020-17026-6

Schrimpe-Rutledge, A. C., Codreanu, S. G., Sherrod, S. D., & McLean, J. A. (2016). Journal of the American Society for Mass Spectrometry, 27(12), 1897–1905. 10.1007/s13361-016-1469-y

Shao, Y., Li, T., Liu, Z., Wang, X., Xu, X., Li, S., Xu, G., & Le, W. (2021). Comprehensive metabolic profiling of Parkinson’s disease by liquid chromatography-mass spectrometry. Molecular Neurodegeneration, 16(1), 4. 10.1186/s13024-021-00425-8

Shen, X., Wang, R., Xiong, X., Yin, Y., Cai, Y., Ma, Z., Liu, N., & Zhu, Z.-J. (2019). Metabolic reaction network-based recursive metabolite annotation for untargeted metabolomics. Nature Communications, 10(1), 1516. 10.1038/s41467-019-09550-x

Southam, A. D., Weber, R. J. M., Engel, J., Jones, M. R., & Viant, M. R. (2017). A complete workflow for high-resolution spectral-stitching nanoelectrospray direct-infusion mass-spectrometry-based metabolomics and lipidomics. Nature Protocols, 12(2), 310–328. 10.1038/nprot.2016.156

Souza, A. L., & Patti, G. J. (2021). A Protocol for Untargeted Metabolomic Analysis: From Sample Preparation to Data Processing. In V. Weissig & M. Edeas (Eds.), Mitochondrial Medicine: Volume 2: Assessing Mitochondria (pp. 357–382). Springer US. 10.1007/978-1-0716-1266-8_27

Sun, Y., Zhang, X., Hang, D., Lau, H. C.-H., Du, J., Liu, C., Xie, M., Pan, Y., Wang, L., Liang, C., Zhou, X., Chen, D., Rong, J., Zhao, Z., Cheung, A. H.-K., Wu, Y., Gou, H., Wong, C. C., Du, L., … Yu, J. (2024). Integrative plasma and fecal metabolomics identify functional metabolites in adenoma-colorectal cancer progression and as early diagnostic biomarkers. Cancer Cell, 42(8), 1386–1400.e8. 10.1016/j.ccell.2024.07.005

Theodoridis, G., Fiehn, O., Holčapek, M., Goodacre, R., Raftery, D., Plumb, R., Ebbels, T. M. D., Witting, M., Gika, H., & Wilson, I. D. (2026). What’s in a name? Metabolite identification: challenges and pitfalls in untargeted metabolomics. Metabolomics, 22(2), 22. 10.1007/s11306-025-02387-0

Vander Heiden, M. G., Cantley, L. C., & Thompson, C. B. (2009). Understanding the Warburg Effect: The Metabolic Requirements of Cell Proliferation. Science, 324(5930), 1029–1033. 10.1126/science.1160809

Wu, X., He, S., Lu, J., Chen, Y., Jiang, X., Wang, X., Zhu, Y., Zhu, Y., Chen, R., Xu, L., & Tang, J. (2025). Untargeted metabolomics and transcriptomics study reveals an activated ferroptosis metabolic spectrum in recurrent spontaneous abortion. Free Radical Biology and Medicine, 239, 145–154. 10.1016/j.freeradbiomed.2025.07.029

Wu, Y., & Wang, Y. (2025). An Improved Metabolomics Workflow Enables Untargeted Data Acquisition and Targeted Data Analysis Using Liquid Chromatography–Mass Spectrometry. Journal of Proteome Research, 24(9), 4734–4743. 10.1021/acs.jproteome.5c00427

