## Supporting Information for "Full Scan enhanced Dynamic Range MS improves metabolite coverage and cancer cell-line discrimination in untargeted metabolomics"

Supplemental Tabel S1: Isolation window boundaries and m/z ranges used for the different Full Scan eDR MS acquisition modes in this study.

| | **Windows** | | | | | | | | | | | | | | | | |
|---|---|---|---|---|---|---|---|---|---|---|---|---|---|---|---|---|---|
| **Scan Mode** | **0** | **1** | **2** | **3** | **4** | **5** | **6** | **7** | **8** | **9** | **10** | **11** | **12** | **13** | **14** | **15** | **16** |
| Full Scan MS | 75 | 1000 | | | | | | | | | | | | | | | |
| Full Scan eDR MS 12 | 75 | 152 | 229 | 306 | 383 | 460 | 537 | 614 | 691 | 768 | 845 | 922 | 1000 | | | | |
| Full Scan eDR MS 14 | 75 | 141 | 207 | 273 | 339 | 405 | 471 | 538 | 604 | 670 | 736 | 802 | 868 | 934 | 1000 | | |
| Full Scan eDR MS 16 | 75 | 133 | 191 | 248 | 306 | 364 | 422 | 480 | 538 | 595 | 653 | 711 | 769 | 827 | 884 | 942 | 1000 |
| Full Scan eDR MS 12 - Custom A | 75 | 125 | 175 | 225 | 275 | 325 | 400 | 475 | 580 | 685 | 790 | 895 | 1000 | | | | |
| Full Scan eDR MS 12 - Custom B | 75 | 95 | 115 | 135 | 155 | 175 | 195 | 215 | 235 | 255 | 275 | 630 | 1000 | | | | |
| Full Scan eDR MS 14 - Custom A | 75 | 125 | 175 | 225 | 275 | 325 | 375 | 445 | 515 | 585 | 668 | 751 | 834 | 914 | 1000 | | |
| Full Scan eDR MS 14 - Custom B | 75 | 100 | 130 | 160 | 190 | 230 | 270 | 310 | 350 | 390 | 430 | 500 | 650 | 800 | 1000 | | |
| Full Scan eDR MS 16 | 75 | 102 | 129 | 156 | 183 | 210 | 237 | 264 | 291 | 318 | 345 | 372 | 399 | 426 | 453 | 727 | 1000 |

**Supplemental Figure S1**

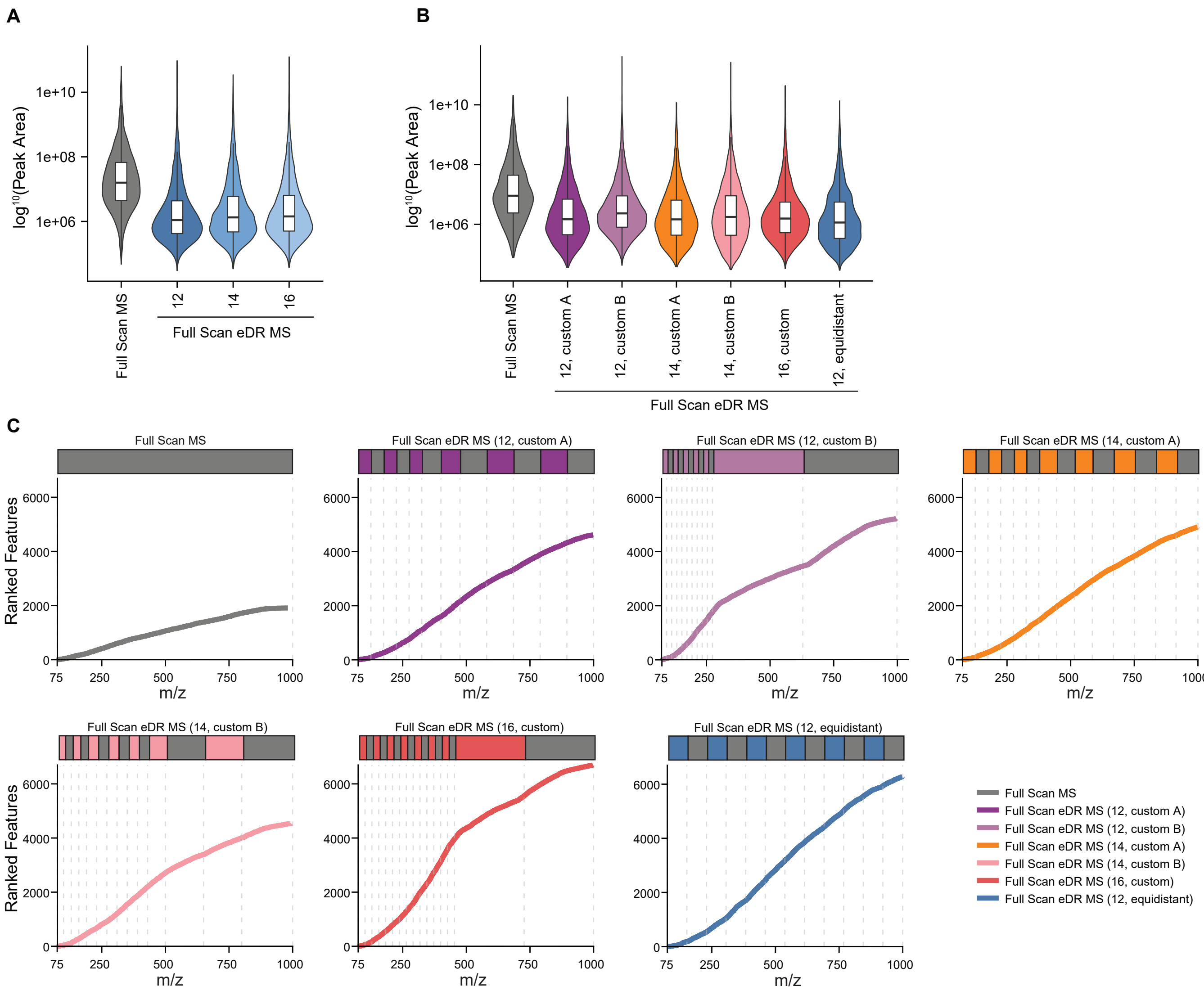


**Supplemental Figure S1**

**A)** Violin plots of compound feature intensities in Full Scan MS or Full Scan eDR MS, using 12, 14, or 16 equidistant isolation windows. **B)** Violin plots of compound feature intensities in Full Scan MS or Full Scan eDR MS for the customized eDR scan mode with 12, 14 or 16 windows with non-uniform isolation windows. **C)** Rank-m/z plots of detected compound features for the customized eDR scan mode with 12, 14 or 16 windows with non-uniform isolation windows, the eDR scanning mode with 12 equidistant windows and Full Scan MS in HepG2 extracts. Grey dashed lines illustrate window distributions as the visualization on top of each graph. Features were identified withCompound Discoverer Software and filtered to retain only those with signal intensities at least threefold higher than blank samples and CV values below 20% across 3 replicates.

**Supplemental Figure 2**

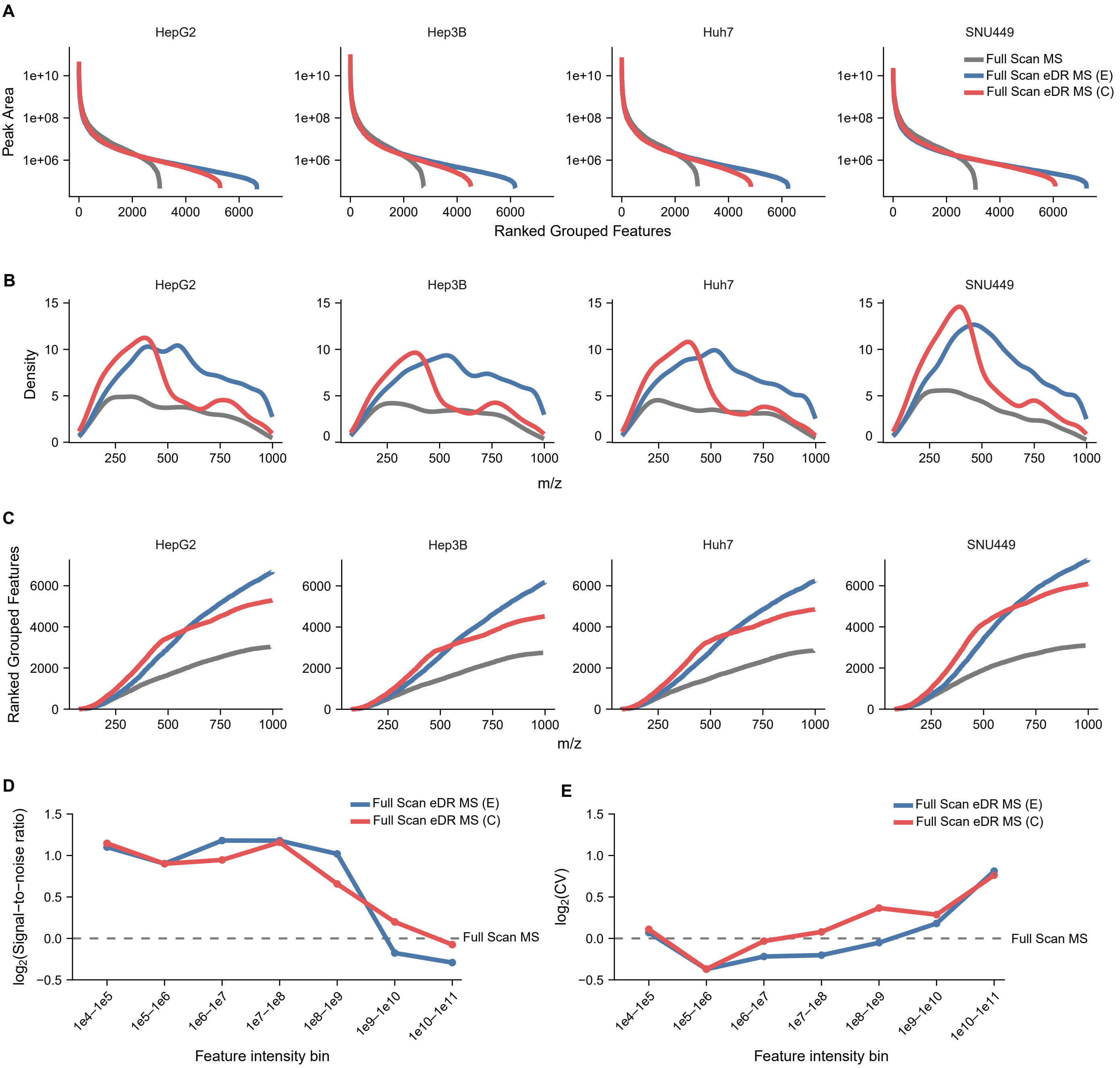


### Supplemental Figure S2

Four human hepatocellular carcinoma (HCC) cell extracts (HepG2, Hep3B, Huh-7, SNU-449) were analyzed in Full Scan MS or Full Scan eDR MS, using 16 windows with either a customized version (C) or with an equidistant distribution (E). Features were identified using Compound Discoverer Software and filtered to retain only those with signal intensities at least threefold higher than blank samples and CV values below 20% across 3 replicates. **A)** Rank-Intensity curve of compound features for all 4 cell lines individually. **B)** Distribution of detected compound features across the m/z range for all 4 cell lines individually. **C)** Rank-m/z curve of compound features for all 4 cell lines individually. **D)** Signal-to-noise ratio (S/N) of detected compound features across intensity bins for the two eDR acquisition modes, relative to Full Scan MS (log2-transformed). Data from all four HCC cell lines were combined. **E)** Coefficients of variation (CV) of detected compound features across intensity bins for the two eDR acquisition modes, relative to Full Scan MS (log2-transformed). Data from all four HCC cell lines were combined.

**Supplemental Figure S3**

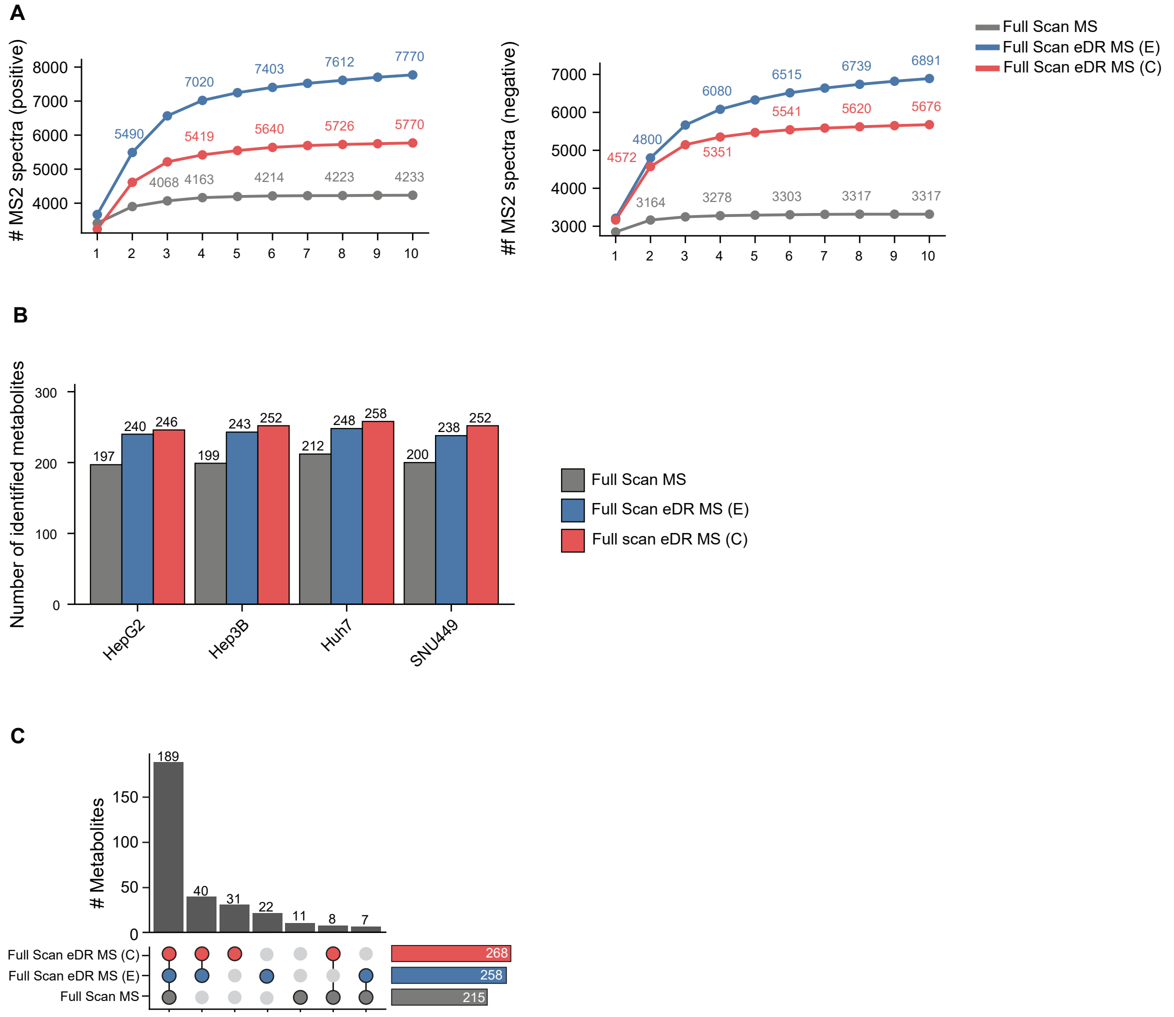


### Supplemental Figure S3

MS2 acquisition and metabolite identification was analyzed in a pooled sample of four human HCC cell extracts (HepG2, Hep3B, Huh-7, SNU-449), using the data-dependent AcquireX deep scan workflow in Full Scan MS or Full Scan eDR MS, using 16 windows with either a customized version (C) or with an equidistant distribution (E). **A)** Total iterative number of newly acquired MS2 spectra per subsequent AcquireX ID run, for each acquisition method across all HCC cell lines in positive and negative mode. **B)** Number of metabolite identifications per acquisition method for each HCC cell line. **C)** UpSet plot showing significantly altered metabolites across cell lines or each acquisition method. Metabolite annotations were generated using mzCloud Spectral Library matching in Compound Discoverer Software with a match factor threshold of 60, and targeted analysis using TraceFinder Software and an in-house library (n=140). Metabolites present in 3 out of 3 replicates per cell line and CV values below 20% were included for downstream analysis.
